# A connection between *Vibrio cholerae* motility and inter-animal transmission

**DOI:** 10.1101/2025.02.12.637895

**Authors:** Ian W. Campbell, Ruchika Dehinwal, Alexander A. Morano, Katherine G. Dailey, Franz G. Zingl, Matthew K. Waldor

**Affiliations:** Division of Infectious Diseases, Brigham & Women’s Hospital, Boston, MA Department of Microbiology, Harvard Medical School, Boston, MA Howard Hughes Medical Institute, Boston, MA, USA

## Abstract

Outbreaks of cholera are caused by the highly transmissive pathogen *Vibrio cholerae*. Here, a transposon screen revealed that inactivation of the *V. cholerae* motility-linked gene *motV* increases infant mouse intestinal colonization. Compared to wild-type *V. cholerae*, a Δ*motV* mutant, which exhibits heightened motility in the form of constitutive straight swimming, localizes to the crypts earlier in infection and over a larger area of the small intestine. Aberrant localization of the mutant was associated with an increased number of *V. cholerae* initiating infection, and elevated pathogen burden, diarrhea, and lethality. Moreover, the deletion of *motV* causes *V. cholerae* to transmit from infected suckling mice to naïve littermates more efficiently. Even in the absence of cholera toxin, the Δ*motV* mutant continues to transmit between animals, although less than in the presence of toxin, indicating that phenotypes other than cholera toxin-driven diarrhea contribute to transmission. Collectively, this work provides experimental evidence linking intra-animal bottlenecks, colonization, and disease to inter-animal transmission.

## Main

Cholera is a severe diarrheal disease caused by *Vibrio cholerae*, a highly motile Gram-negative rod ^1^. Individuals become exposed to the pathogen by ingesting contaminated water or food or through close person-to-person spread ^2–4^. After ingestion, the pathogen colonizes and rapidly proliferates in the small intestine (SI). During its growth in the SI, *V*. *cholerae* secretes cholera toxin, an AB_5_-type protein toxin whose activities largely account for the secretory diarrhea characteristic of cholera ^5^.

Orogastric inoculation of 3-5-day-old infant mice with toxigenic *V*. *cholerae* has been used to model cholera for over five decades ^6–8^. The disease observed in infant mice mirrors many aspects of human infection, including toxin-dependent diarrheal disease ^9^. Studies in this model have yielded many critical insights into the genes and processes that contribute to *V. cholerae* pathogenicity ^10,11^. For example, toxin co-regulated pilus (TCP), the pathogen’s signature colonization factor, was initially identified in infant mice^12^ and was subsequently demonstrated to be critical for human infection ^13^.

Infant mouse studies also revealed that *V. cholerae* motility and chemotaxis play important and often opposite roles in intestinal colonization. The pathogen has a single polar sheathed flagellum whose counterclockwise rotation causes straight swimming. Chemotactic input causes random reorientation by briefly reversing the spin of the flagellum to clockwise rotation ^14^. Nonmotile mutants lacking a flagellum or with an inactive flagellum (e.g., Δ*motAB*) exhibit marked defects in colonizing infant mice ^15,16^. Conversely, chemotaxis mutants with an active flagellum often exhibit heightened colonization ^17^. For example, *V. cholerae* chemotaxis mutants that cannot change the direction of flagellar rotation from counterclockwise to clockwise (e.g., Δ*cheY*) exhibit longer stretches of straight swimming and have increased intestinal colonization, particularly in the proximal SI ^18^.

Cholera epidemics are often characterized as ‘explosive’ because of the rapid spread of the disease to exposed individuals ^1^. Animal colonization models have elucidated many aspects of pathogenesis that could contribute to the characteristic transmissibility of *V. cholerae*, including pathogen colonization factors that facilitate replication within the intestine, a host-associated increase in pathogen infectivity (aka hyperinfectivity), and the role of virulence genes in diarrhea ^10^. However, the mechanisms resulting in pathogen transmission are usually not captured in animal models because infections are initiated by directly inoculating cultured bacteria into the stomach, limiting our knowledge of the factors that govern *V. cholerae* transmissibility. Here, we adapted the infant mouse model to quantify transmission between pups and leveraged a *V. cholerae* motility mutant to understand how events within the intestine feed-forward to govern transmission.

### Genome-scale screen to identify *V. cholerae* genes modifying infant mouse colonization

Transposon insertion site sequencing (Tn-seq) is a powerful approach for genome-scale identification of bacterial genes that modify growth in diverse conditions. In animal-based studies, Tn-seq screens are generally carried out to identify mutants that fail to colonize, suggesting that the transposon-disrupted gene promotes bacterial growth in the host. However, Tn-seq has not yet been used to define the genes required for *V. cholerae* colonization of infant mice.

Colonization bottlenecks can render Tn-seq studies uninterpretable because the bottleneck causes a stochastic loss of mutants from the population, which can obscure the identity of mutants lost due to selective pressure in the host. However, recent studies measuring the bottleneck restricting the *V. cholerae* population during colonization of infant mice suggest that Tn-seq is possible in this model; using genomically-barcoded *V. cholerae*, Gillman et al. (2021) demonstrated that when given a high dose (10^8^ colony forming units, CFU), ∼10^5^ unique bacterial cells survive the bottleneck and expand in the SI of infant CD1 mice ^19^. High-density Tn-seq libraries created with a mariner transposon generally contain ∼10^5^ unique mutants, approximately the same number of cells that survive the bottleneck. We used this insight to carry out the first Tn-seq screen for *V. cholerae* genes that modify growth in the infant mouse intestine.

We orally inoculated ∼5x10^7^ CFU of a dense transposon library containing ∼1.1x10^5^ unique mutants created in a 2010 *V. cholerae* clinical isolate from Haiti ^20^ into P4 CD1 mice. 18-hours after inoculation, we recovered between 1.7 to 2.5 x10^4^ of 1.1x10^5^ unique mutants from the SI of 3 pups (Fig S1A), indicating that the bottleneck caused a substantial loss of mutant diversity. To increase the recoverable mutants, we pooled the SIs of 10 P4 mice (one litter) and recovered 3.3x10^4^ unique mutants, covering ∼35% of the average gene’s potential mutagenesis sites (Fig S1A-B). This degree of mutant loss can be accommodated by our analytical pipeline, which uses simulation-based normalization to model and compensate for the random loss of diversity caused by population bottlenecks ^21,22^.

Using this approach, we identified the genes that promote *V. cholerae* survival in the infant mouse SI (Table S1). Transposon insertions in genes that facilitate growth in the murine intestine but not when cultured in LB media are found on the left side of the volcano plot in Fig 1A. The “hits” on this side of the plot validate this application of Tn-seq. Transposon insertions in many genes previously demonstrated to be critical for intestinal colonization in this model, such as genes required for the biogenesis of the type IV pilus TCP (shown in orange in Fig 1A-B), had markedly reduced abundance *in vivo*. Within the *tcp* locus, the reduction in the abundance of insertions in genes such as *tcpA* ^23^, the major subunit of the pilus, and *tcpF*, a secreted and essential colonization factor ^24^, was greater than 1,000-fold; in contrast transposon insertions were not depleted in *tcpI* during colonization, a gene that is not required for colonization and that has been linked to reduced TcpA expression ^25^, illustrating the specificity of the Tn-seq screen.

**Figure 1.**
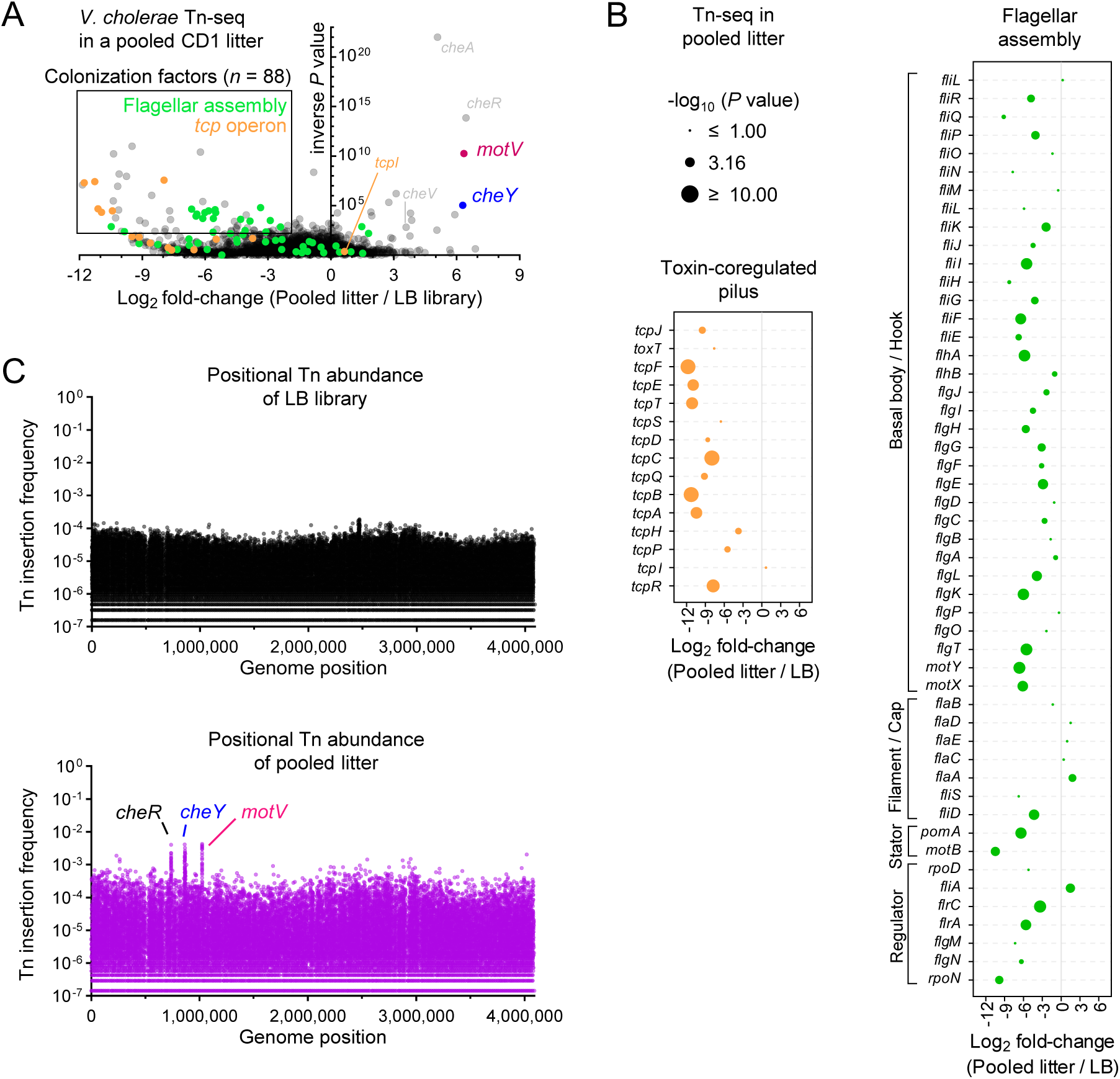
Tn-seq screen reveals that disruption of *motV* increased *V. cholerae* infant mouse colonization. 10 CD1 pups were intragastrically inoculated with ∼5x10^7^ CFU of a mariner transposon library. 18-hours later, the SIs were homogenized and pooled, and after overnight growth on LB plates, transposon insertion sites were determined by sequencing (pooled litter). For comparison, the same library was passaged overnight on LB plates (LB library). **A,** Volcano plot comparing the fold-change in gene insertion frequency between the pooled litter and LB library. Colonization factors are genes with *P* value <0.01 and log_2_ fold-change <-2. Data for the *tcp* operon (toxin-coregulated pilus) and flagellar assembly genes (categorized from KEGG database vch02040) are expanded in **B**. **C,** Transposon insertion frequency across the genome highlighting increased insertion frequency in *cheR, cheY,* and *motV* from the pooled intestinal samples.

Flagellar-based motility is generally thought to promote *V. cholerae* intestinal colonization ^14^, and insertions in many of the genes linked to the biogenesis and spinning of the pathogen’s flagellum had reduced abundance in animal samples relative to growth in LB (Fig 1A-B). In contrast, there was no reduction in insertions in the flagellin genes *flaABCDE*, which are the building blocks of the flagellar filament. The lack of a phenotype associated with disruption of *flaABCDE* may be due to functional redundancy between the flagellins. However, unlike *flaBCDE, flaA* was previously determined to be necessary and sufficient for motility in another *V. cholerae* strain ^26^, and it is unclear why *flaA* is not required for colonization here.

Intriguingly, several genes were also identified on the right side of the volcano plot (Fig 1A). Insertions in these genes were more abundant *in vivo* than *in vitro*, suggesting that these genes antagonize *V. cholerae* growth in the intestine. Within the initial LB library, no individual mutant was represented by more than 0.02% of sequencing reads. Mutants that became more abundant in the animals were visualizable as peaks in transposon abundance, with individual mutants comprising >0.4% of reads (Fig 1C). Four of these genes, *cheA*, *cheR*, *cheV* and *cheY* (Fig 1A), are linked to chemotaxis, which was previously demonstrated to limit colonization ^14^. *motV* insertions were among the most enriched *in vivo* (Fig 1A, 1C). The molecular function of MotV is unknown, but it was previously demonstrated to control motility and associate with HubP ^27^, the *V. cholerae* cell pole organizer ^28^. Without *motV*, *V. cholerae* cannot reverse directions (tumble). Cells lacking *motV* have increased motility in liquid, constitutively swimming straight at elevated speeds. Conversely, cells lacking *motV* cannot swim in soft agar, which requires reversing direction ^27^. However, MotV’s role in *V. cholerae* infection has not been studied.

### Deletion of *motV* improves *V. cholerae* intestinal colonization

To corroborate that *motV* antagonizes *V. cholerae* colonization, we created an in-frame *motV* deletion mutant (Δ*motV)*. As expected, the Δ*motV* strain had severely impaired motility in soft agar, like that observed in Δ*cheY* mutants (Fig S2A-B), and the provision of a plasmid-borne copy of *motV* (p*motV*) restored soft agar motility (Fig S2C). In infant mice, the number of Δ*motV* CFU recovered from the SI exceeded the CFU recovered from animals infected with wild-type (WT) *V. cholerae* and was similar to the CFU found with a Δ*cheY* mutant (Fig 2A), which is known to increase colonization ^16^. Further analyses found that the heightened colonization of the Δ*motV* strain was particularly prominent in the proximal SI, where Δ*motV* CFU exceeded those of the WT by ∼13-fold (Fig 2B). Ordinarily, ∼10x more WT V. cholerae CFU are recovered from the distal SI than the proximal SI. However, similar CFU of the Δ*motV* mutant strain were recovered from the two parts of the intestine. The elevated capacity of the Δ*motV* mutant to colonize the proximal SI was reduced by the provision of p*motV* (Fig S2D), suggesting that this Δ*motV* phenotype is specifically caused by the deletion of *motV*. Moreover, the heightened colonization capacity of the Δ*motV* strain was confirmed in competition assays, where *lacZ*^+^ Δ*motV* outcompeted a *lacZ*^-^ WT strain 13-fold in the proximal SI and 4-fold in the distal SI (Fig 2C).

**Figure 2.**
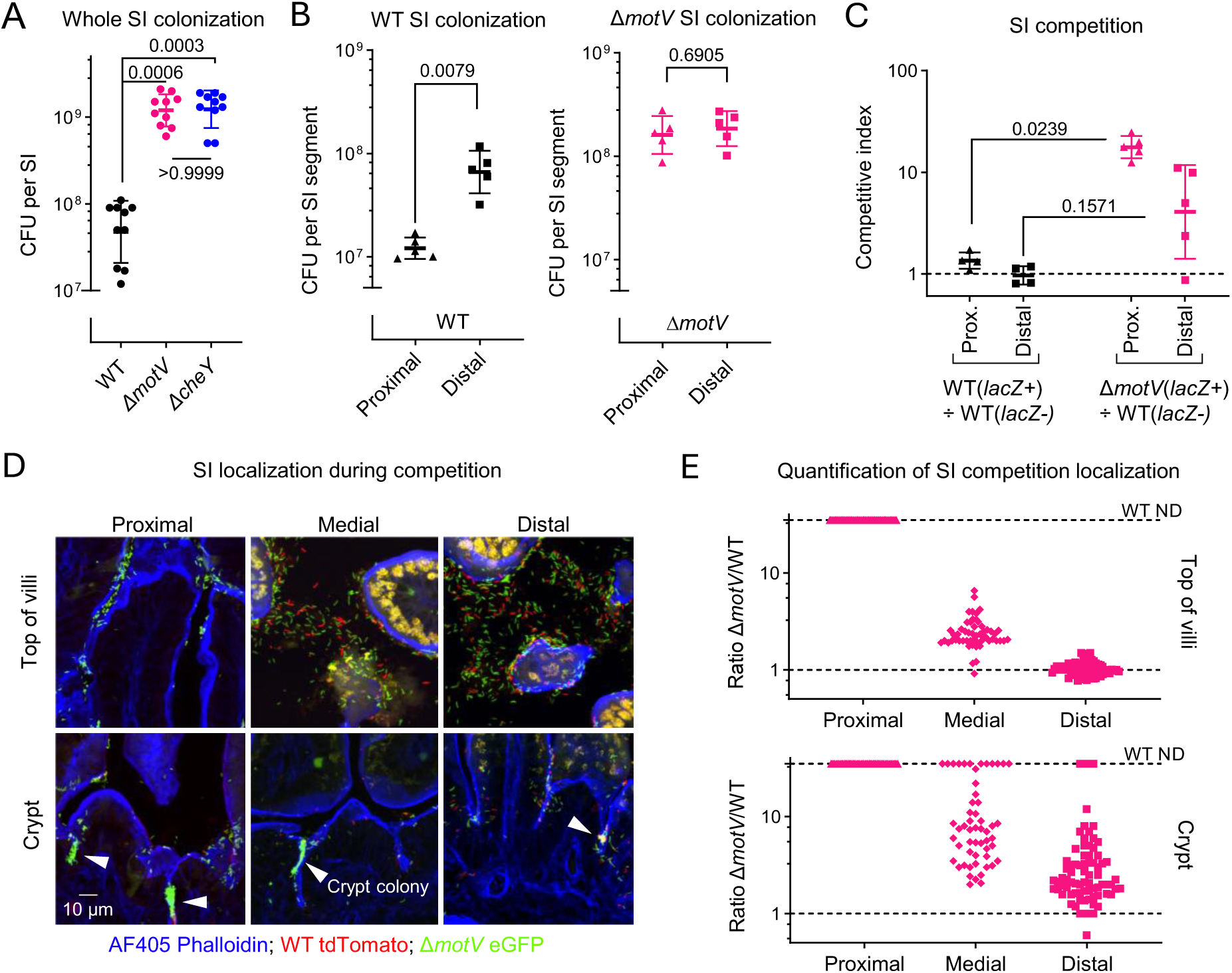
Deletion of *motV* increased *V. cholerae* SI colonization, especially in the proximal SI and within the crypts. CD1 pups were intragastrically inoculated with ∼3 x 10^6^ CFU of the indicated *V. cholerae* strains, and 18-hours later the whole intestine (**A**) or the indicated SI segments (**B**) were plated on selective media to determine *V. cholerae* colony-forming units (CFU). **C,** CD1 pups were infected with ∼2 x 10^6^ CFU of a ∼1:1 ratio of *lacZ^+^* to *lacZ^-^ V. cholerae*, and the competitive index was calculated as the ratio of *lacZ^+^* to *lacZ^-^* CFU in the indicated SI segment 18-hours after inoculation divided by the ratio of *lacZ^+^* to *lacZ^-^*CFU in the inoculum. **A** and **C,** Kruskal-Wallis test with Dunn’s multiple comparison correction. **B,** Mann-Whitney test. **A-C,** Geometric mean and standard deviation. **D,** Images of 10 µm cryosections of the indicated SI sections 18-hours after inoculation with ∼2 x 10^6^ CFU of a ∼1:1 ratio of WT *lacZ::tdTomato V. cholerae* (red) and Δ*motV lacZ::eGFP V. cholerae* (green). Sections were stained with AF405-conjugated phalloidin (blue) and imaged on a spinning disk confocal microscope. Arrowheads show *V. cholerae* microcolonies. **E,** Quantification of the ratio of Δ*motV*:WT *V. cholerae* cells at the top(s) of the villi and in the crypts of the SI. When WT was not detected (ND), the ratio was given a value of 35 as an upper limit. *V. cholerae* were differentiated from background fluorescence by comma/rod morphology.

A 1:1 ratio of eGFP-tagged Δ*motV* and tdTomato-tagged WT *V. cholerae* were co-inoculated into infant mice to determine if *motV* deletion changed the pathogen’s localization within the SI. Notably, previous work found that fluorophore expression does not measurably alter *V. cholerae* fitness ^29^. There was a much greater abundance of the Δ*motV* strain compared to the WT strain, particularly in the proximal SI, where it was difficult to detect the WT strain (Fig 2D-E). Besides corroborating the competitive advantage of the Δ*motV* mutant along the proximal to distal axis of the SI, the fluorescence microscopy also revealed a differential localization of the two strains along the crypt-villus axis. In all segments of the SI, the Δ*motV* mutant had an enhanced capacity to access the crypts and form microcolonies (Fig 2D-E; Fig S3). Thus, the absence of *motV* enhances colonization at least in part by facilitating the pathogen’s entry into the SI crypts, especially in the proximal SI.

### The Δ*motV* mutant localizes early to the SI crypts

To gain insight into how deletion of m*otV* augments *V. cholerae* intestinal colonization, we compared the WT and mutant populations throughout the course of infection. There was a pronounced collapse in the size of the WT population over the first 3-hours of infection, especially in the proximal SI (Fig 3A), as previously described ^30^. After the first 3-hours, the WT population expanded in the proximal, medial, and distal SI. Notably, the size of the Δ*motV* mutant population was much less constricted over the first 3-hours, particularly in the proximal SI. Subsequently, the Δ*motV* population remained more numerous than the WT population in the proximal SI from 3-24 hours, while the populations approached parity in the medial and distal SI at later timepoints.

**Figure 3.**
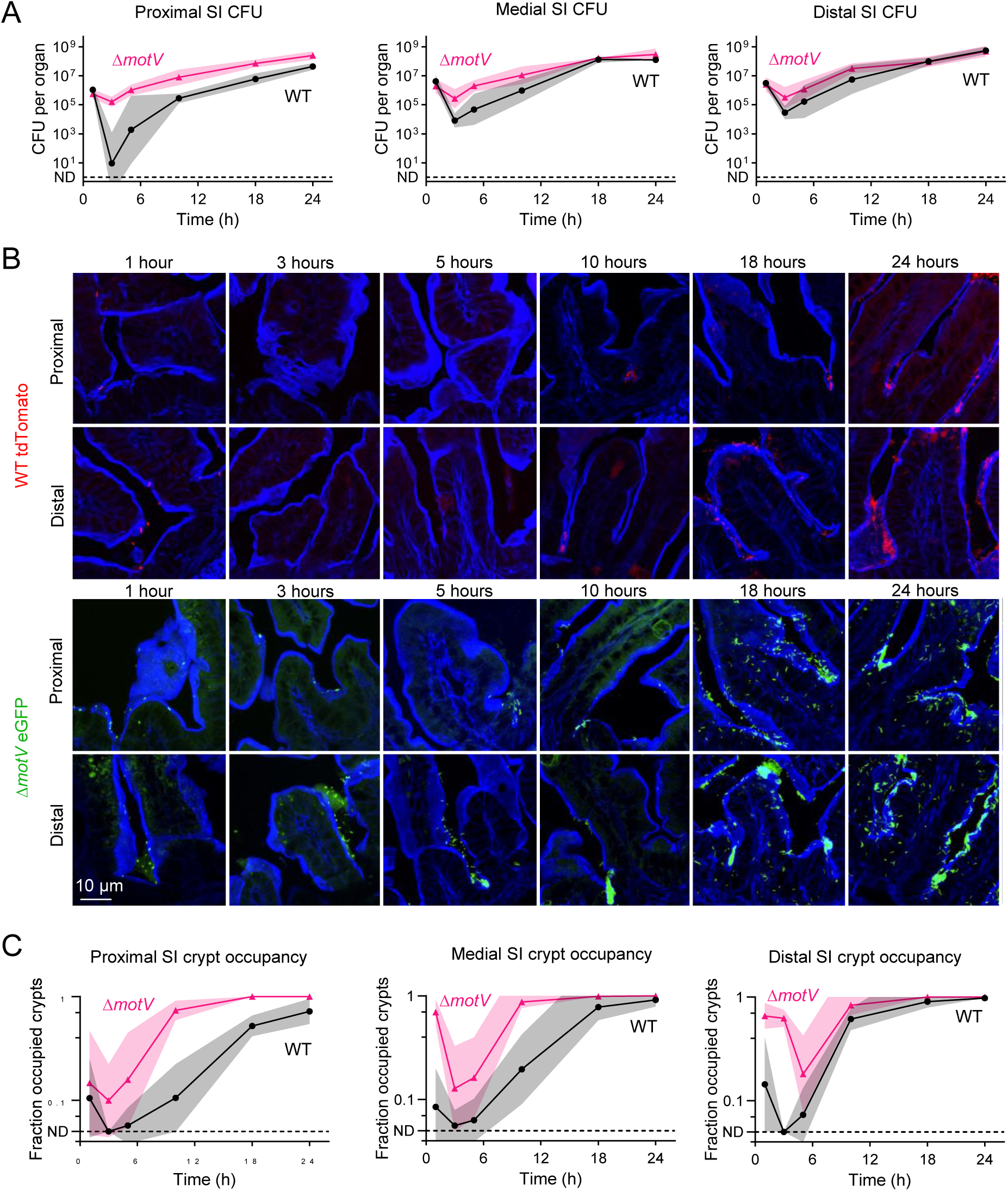
The Δ*motV* mutant localizes in the SI crypts earlier during infection than WT *V. cholerae*. CD1 pups were intragastrically inoculated with ∼2 x 10^6^ CFU of either WT *lacZ::tdTomato V. cholerae* (red) or Δ*motV lacZ::eGFP V. cholerae* (green). CFU (**A**) and cryosections were collected over 24-hours from the indicated SI sections. Sections were stained with AF405-conjugated phalloidin (blue) and imaged on a spinning disk confocal microscope (**B**). **C,** Quantification of the number of crypts per field occupied by at least one *V. cholerae* in the proximal, medial, and distal SI. ND, *V. cholerae* not detected in any field. 4 animals per timepoint per strain. **A, C,** Geometric mean and standard deviation.

We hypothesized that the constitutive straight swimming motility phenotype of the Δ*motV* mutant might contribute to the pathogen’s capacity to persist in the proximal SI. To test this hypothesis, we used fluorophore-labeled *V. cholerae* to measure the pathogen’s localization throughout the course of infection (Fig 3B). Within the first hour, when the total burden was similar for both strains (Fig 3A), it was already apparent that the Δ*motV* mutant was more numerous in the intervillous crypts. We quantified the fraction of crypts per field that were occupied by at least one *V. cholerae* at each timepoint (Fig 3C). Throughout the infection, a greater fraction of crypts in the proximal small intestine were occupied by Δ*motV* than WT. Greater crypt occupancy by the Δ*motV* mutant was also observed in the medial and distal SI for most of the infection, with wild-type reaching parity at 24-hours. The increased penetration of the Δ*motV* mutant into the crypts may result from its constitutive straight swimming and suggests that the motility behavior of the wild-type strain is insufficient to efficiently reach the crypt niche, particularly early in infection.

### The Δ*motV* mutant circumvents the host bottleneck

We hypothesized that the failure of WT *V. cholerae* to reach the crypts and persist in the proximal SI during the first hours following inoculation may contribute to the initial collapse of the population and account for at least a portion of the WT colonization bottleneck. To directly characterize how deletion of *motV* impacts the *V. cholerae* colonization bottleneck, we used ‘STAMPR’, a lineage tracing method that tracks bacterial population dynamics during infection using otherwise genetically identical cells identified by unique DNA barcodes ^31^. Comparing the frequency and diversity of barcodes in the inoculum to samples from host tissues enables measurement of the number of unique bacterial cells (founding population) from which the observed population originated (Fig 4A). The change in the number of cells from the inoculum to the founding population quantifies the infection bottleneck - the set of factors (e.g., physical barriers, immune processes, and the microbiota) that limit the capacity of a bacterial population to establish infection, with smaller founding populations reflecting tighter (more restrictive) infection bottlenecks.

**Figure 4.**
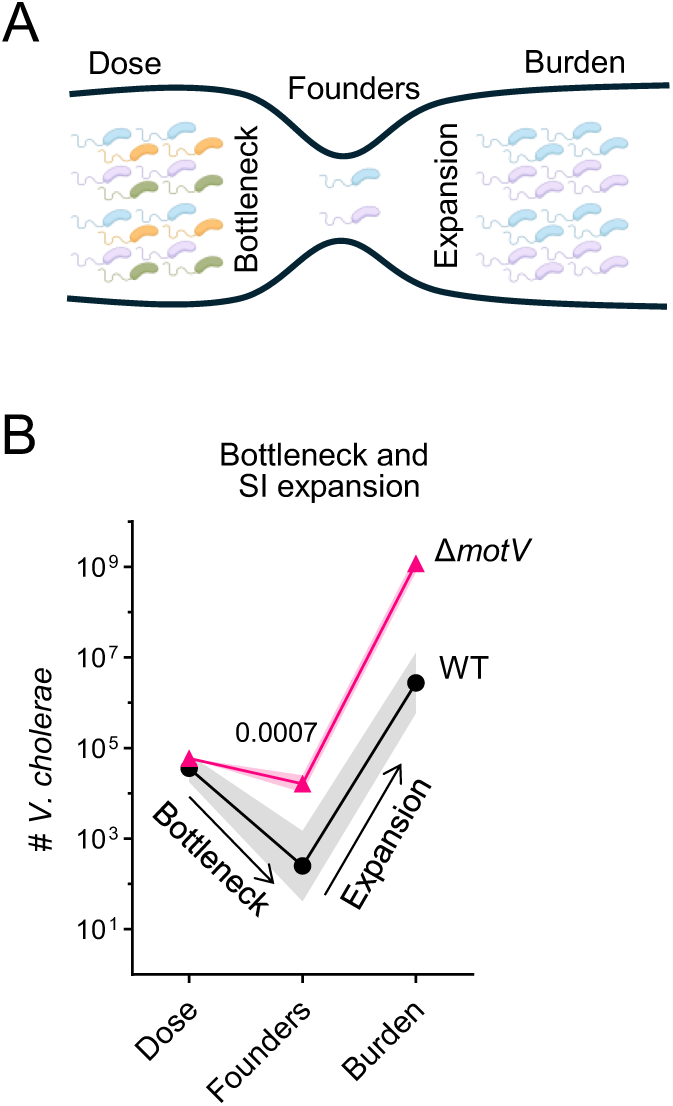
*V. cholerae* lacking *motV* evaded the colonization bottleneck. **A,** Diagram of bacterial population dynamics during infection. The dose of the inoculum and the burden of the pathogen in tissue is enumerated by plating for CFU on selective media. The founding population is the number of unique cells from which the observed population originated and is measured by the loss of barcode diversity (represented as colors) between the inoculum and the observed population. Founders are calculated by STAMPR and expressed as Nr. The bottleneck is the loss of cells between the dose and founders, and net expansion is the gain between founders and burden. **B,** Dose, founders, and burden of WT and Δ*motV V. cholerae* in the SI of CD1 pups 18-hours post-inoculation. WT, 5 pups; Δ*motV*, 10 pups. Mann-Whitney test. Geometric mean and standard deviation.

The increased burden of the Δ*motV* mutant in the SI compared to WT could be due to the strain undergoing a less restrictive bottleneck, leading to a larger founding population and/or more net replication. To distinguish between these possibilities and quantify the Δ*motV* bottleneck, we used STAMPR to compare the population dynamics of barcoded WT and Δ*motV* bacteria in P5 mice. When administered similar-sized inoculums, the founding population of the Δ*motV* strain in the SI was ∼100-fold greater than that measured for the WT strain (Fig 4B), indicating that the mutant experienced a much less restrictive bottleneck. Surprisingly, the Δ*motV* strain only experienced a ∼2-fold bottleneck, with half of the Δ*motV* cells in the inoculum surviving to become the founding population (Fig. 4B). Thus, the absence of *motV* allows *V. cholerae* to evade the factors that underlie the infection bottleneck. We propose that the ability of the Δ*motV* mutant to evade host bottlenecks is caused by its constitutive straight swimming phenotype, which allows cells to persist in the proximal SI and invade deeper into the intestinal crypts (Fig 3), forming replicative microcolonies (Fig 2D).

### Deletion of *motV* increases *V. cholerae* virulence

We observed that the infant mouse bedding was more stained with diarrheal discharge in animals infected with the Δ*motV* mutant than those infected with the WT strain (Fig S4), suggesting that the *ΔmotV* mutant induced more diarrhea in addition to increasing colonization. Indeed, the weight of the bedding, a measure of the diarrheal discharge, was greater in the *ΔmotV* group than WT (Fig 5A). Furthermore, the mice infected with the mutant strain had more weight loss and a greater ratio of SI/body weight (a measure of fluid accumulation) than mice infected with the WT strain (Fig 5A). Although the Δ*cheY* mutant colonized at least as well as the Δ*motV* mutant (Fig 2A), it did not lead to increased diarrheal discharge or weight loss (Fig 5A). Thus, the increased colonization of the Δ*motV* strain likely does not entirely account for its elevated diarrheaogenicity. The elevated virulence of the Δ*motV* strain was also apparent in survival studies^32,33^, where pups were returned to dams, and morbidity was monitored until a predetermined 30-hour endpoint. The Δ*motV* mutant resulted in mortality in infected animals more rapidly than either the WT or Δ*cheY* strains (Fig 5B).

**Figure 5.**
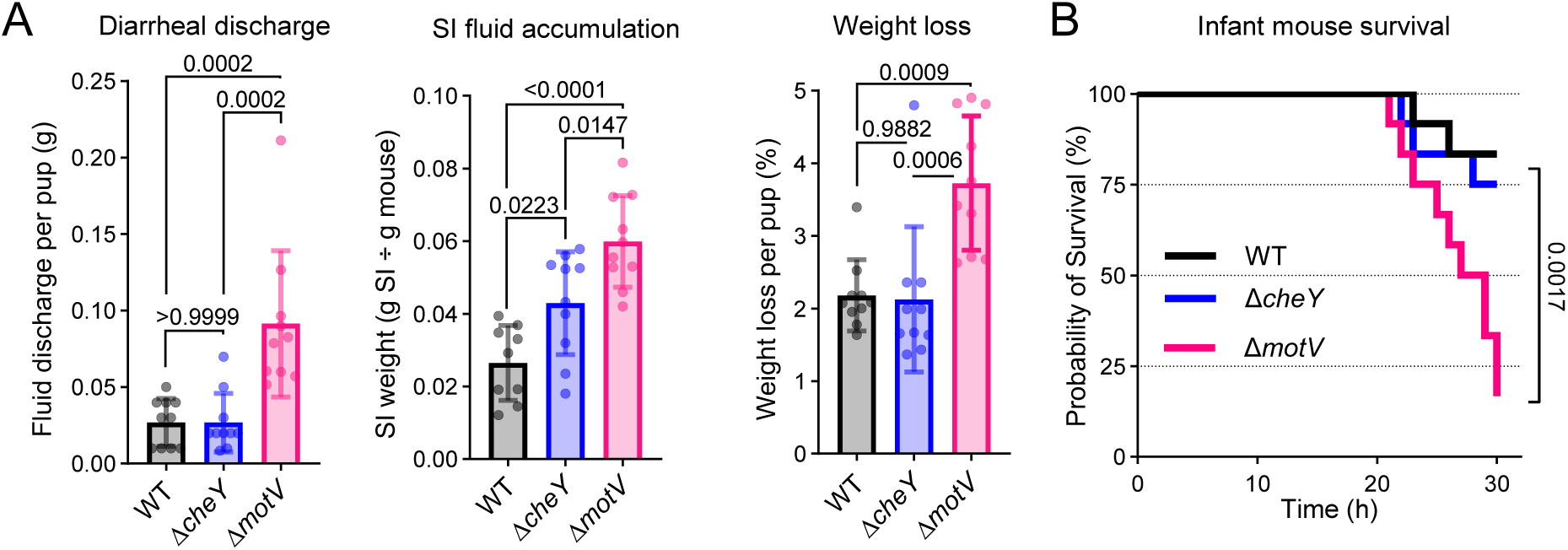
Deletion of *motV* increased *V. cholerae* diarrheaogenicity. **A,** CD1 pups were intragastrically inoculated with the indicated *V. cholerae* strains and individually housed for 18-hours. The bedding was weighed prior to and after infection to determine fluid discharge. Fluid accumulation in the SI 18-hours post inoculation was measured by comparing the weight of the animal to the weight of the SI. Weight loss was determined by comparing animal weight before and after infection. Mean and standard deviation. One-way ANOVA with Turkey’s multiple comparison correction. **B,** CD1 pups were intragastrically inoculated with the indicated *V. cholerae* strains, returned to dams for care, and monitored until the predetermined 30-hour endpoint. Survival kinetics with Mantel-Cox test.

The principal cause of *V. cholerae*-associated diarrhea is cholera toxin ^5^. To test whether deletion of *motV* increases diarrheaogenicity by causing cells to produce more toxin, we measured toxin production in laboratory conditions that induce cholera toxin synthesis (AKI) ^34,35^. The WT, Δ*motV*, and Δ*cheY* strains produced similar quantities of cholera toxin in these conditions (Fig S5). Since both the Δ*cheY* and Δ*motV* mutants exhibit defective motility characterized by an inability to tumble and swim in soft agar^27^ (Fig S2), increased burden during infection (Fig 2A), and similar production of cholera toxin in culture, the cause of the increased virulence of the Δ*motV* mutant remains unclear.

### The Δ*motV* mutant is hyper-transmissible

Many of the phenotypes of the Δ*motV* mutant are associated with transmission, including bottleneck evasion, replication to high burdens in the intestine, and shedding into the environment in diarrhea. To experimentally test whether these characteristics feed-forward to increase spread between animals, we devised a method to quantify *V. cholerae* transmissibility (Fig 6A). In this system, post-natal day 4 mice from several litters were randomized into new litters to minimize litter bias. The next day, ∼1/3 of the mice in each new litter were infected with the indicated *V. cholerae* strain; these infected ‘seed’ mice were then mixed with approximately twice as many uninfected ‘contact’ mice (the remaining 2/3 of the new litter) and returned to foster dams in individual cages for 20 hours (Fig 6A). At that point, the number of CFU in the SI of seed and contact groups was quantified to determine the number of infected contacts and the robustness of their colonization.

**Figure 6.**
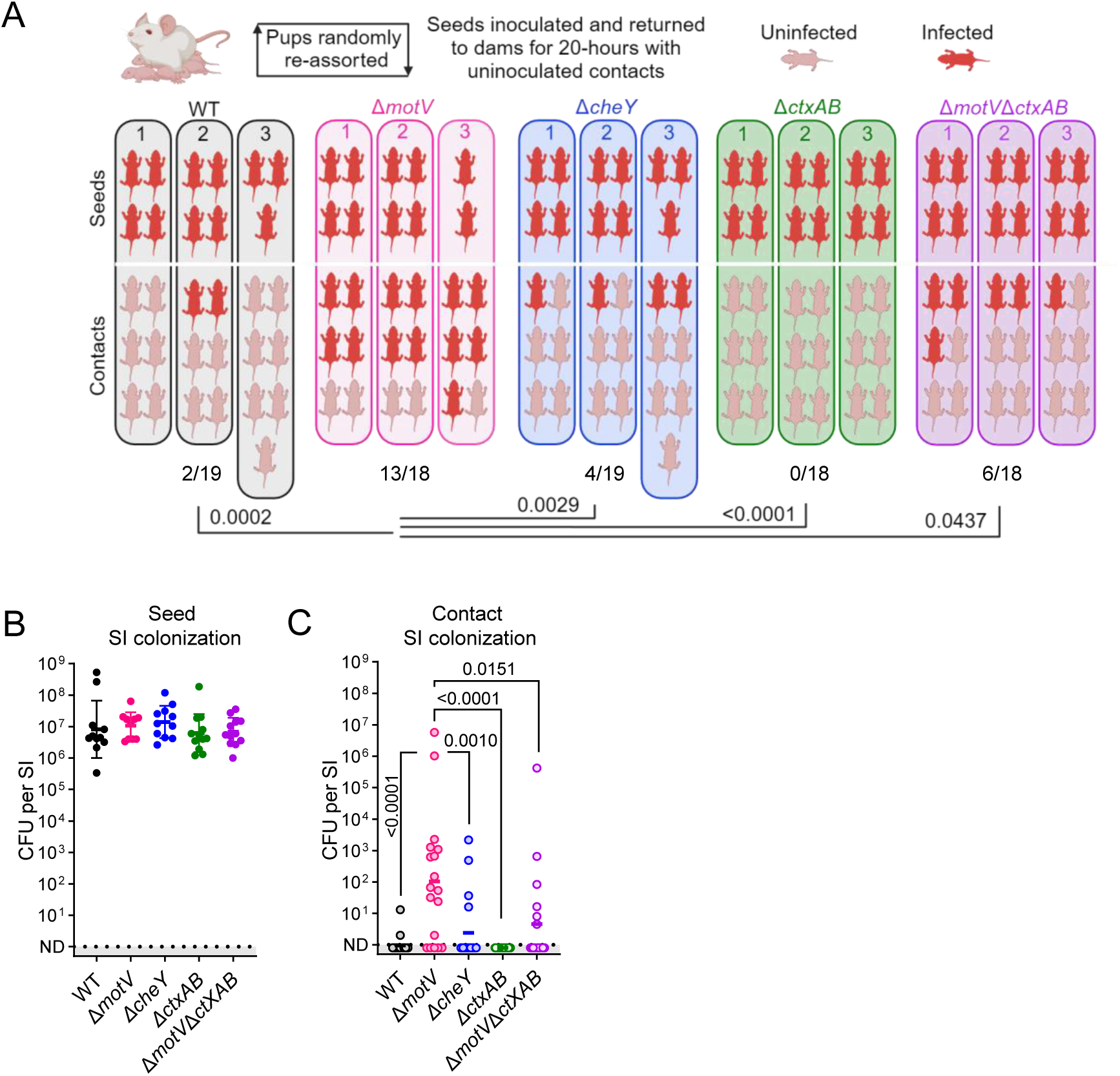
Deletion of *motV* increased inter-animal transmission. Transmission assays were performed by randomly reassorting CD1 pups, challenging ∼1/3 of the litter with the indicated strain of *V. cholerae* (seeds), and returning them to foster-dams with naïve littermates (contacts). 20-hours later, transmission was determined by enumerating CFU in the SI. Experimental groups: WT, Δ*motV,* and Δ*cheY;* Δ*ctxAB* and Δ*motV* Δ*ctxAB*. **A,** Experimental results divided by litter. Uninfected contacts, 0 CFU detected; infected, ≥1 CFU identified. **B,** Seed SI CFU determined 20-hours post inoculation. Geometric mean and standard deviation. **C,** Contact SI colonization determined after 20-hours of cohousing with seed animals. Geometric mean. Kruskal-Wallis test with Dunn’s multiple comparison correction.

Across three independent trials, seed animals infected with the Δ*motV* mutant were more transmissive than WT (Fisher’s exact test *P* value 0.0002); 13/18 (72%) of contacts of Δ*motV* seeds were infected, whereas 2/19 (11%) of contacts of WT seeds were infected (Fig 6A). Additionally, contact animals that were infected by Δ*motV* seed animals tended to have greater intestinal colonization than contacts of the WT seed animals (Fig 6C). Thus, the Δ*motV* mutant is ‘hyper-transmissible’, indicating that MotV impedes *V. cholerae* transmission as well as colonization. To our knowledge, the Δ*motV* strain represents the first description of a hyper-transmissible *V. cholerae* mutant.

We leveraged the Δ*cheY* mutant to test whether the hyper-transmissibility of the Δ*motV* mutant is solely attributable to its increased intestinal colonization. Like *motV* mutants, strains lacking *cheY* exhibit increased colonization (Fig 2A and ^15^^,16,18^). However, the Δ*cheY* mutant was not hyper-transmissible in this assay, transmitting to fewer contacts (4/19; 21%) than Δ*motV* (Fig 6A). These results suggest that the increased colonization of Δ*motV* does not fully account for its enhanced transmission.

Since the hyper-transmissibility of the Δ*motV* mutant is not solely attributable to increased colonization, we hypothesized that increased transmission could be caused by increased shedding of the pathogen in the more abundant diarrhea of mice infected by the mutant, a phenotype not shared with Δ*cheY*. To test this hypothesis, we used a Δ*ctxAB* mutant that does not produce cholera toxin, the principal cause of *V. cholerae*-associated diarrhea ^5^. As expected, mice infected with strains lacking *ctxAB* or lacking both *ctxAB* and *motV* had less fluid accumulate in their SI than mice infected with the Δ*motV* mutant (Fig S6). There was no transmission from seed animals infected with a strain lacking *ctxAB* (0/18; 0%; Fig 6A), suggesting that cholera toxin promotes the transmission of WT *V. cholerae* in this system. Furthermore, seed animals infected with the Δ*motV* Δ*ctxAB* double mutant were less transmissive than seed mice infected with *V. cholerae* lacking *motV* alone (Fig 6A, C), suggesting that cholera toxin-driven diarrhea contributes to transmission of the Δ*motV* mutant.

Nevertheless, cholera toxin was not required for the Δ*motV* mutant to transmit, as the Δ*motV* Δ*ctxAB* double mutant transmitted to 6/18 (33%) of contacts (Fig 6A). Notably, since seed animals infected with the Δ*motV* Δ*ctxAB* double mutant were more transmissive than seeds infected with a Δ*ctxAB* single mutant, a portion of the Δ*motV’s* hyper-transmissibility is independent of cholera toxin-induced diarrhea.

## Discussion

A hallmark of infectious agents is their capacity for transmission between hosts. However, there is a relative paucity of experimental models to measure pathogen transmissibility, limiting our knowledge of the genes and processes involved in transmission. Here, we found that suckling mice infected with *V. cholerae* could transmit the pathogen to uninfected littermates, providing an experimental model of transmission. We leveraged this model to characterize how the motility-linked gene *motV* modifies *V. cholerae* transmission. The Δ*motV* mutant localized to the intestinal crypts earlier and over a larger area of the small intestine than wild-type *V. cholerae*. This change in localization was associated with 3 phenotypes that likely drive the hyper-transmissibility of the Δ*motV* mutant: (1) evasion of the host bottleneck, (2) elevated SI colonization, and (3) increased diarrheaogenicity.

Our findings suggest a model to describe how the constitutive straight swimming phenotype of the Δ*motV* mutant may result in hyper-transmissibility (Fig 7). While WT chemotaxis inefficiently localizes *V. cholerae* to the crypt niche during the first hours following inoculation (Fig 3B-C), deletion of *motV* enables earlier and more efficient occupancy of the intestinal crypts, especially in the proximal SI. We propose that the aberrant pattern of localization of the Δ*motV* mutant underlies many of its other phenotypes. Thus, early crypt occupancy may enable the Δ*motV* mutant to escape peristaltic flow and linger longer in the proximal SI, potentially explaining why the Δ*motV* mutant experiences a less restrictive bottleneck. Furthermore, occupying more of the SI likely enables the Δ*motV* strain to reach a higher burden, and the higher burden combined with proximity to the intestinal epithelium likely contributes to greater intoxication of the host and increased pathogen shedding. Moreover, shed Δ*motV V. cholerae* avoids the bottleneck and colonizes the next host at a lower dose. Collectively, aberrant motility leads to more cells from the inoculum initiating infection, elevated burden, and increased diarrhea, which together feed-forward to heighten transmission. Our findings illuminate the processes that are likely exploited by many enteric pathogens for their characteristic transmission: (1) evasion of host barriers, (2) replication to high burdens in infected tissues, and (3) dissemination in diarrhea, and demonstrate how intra-animal pathogenesis results in inter-animal transmission.

**Figure 7.**
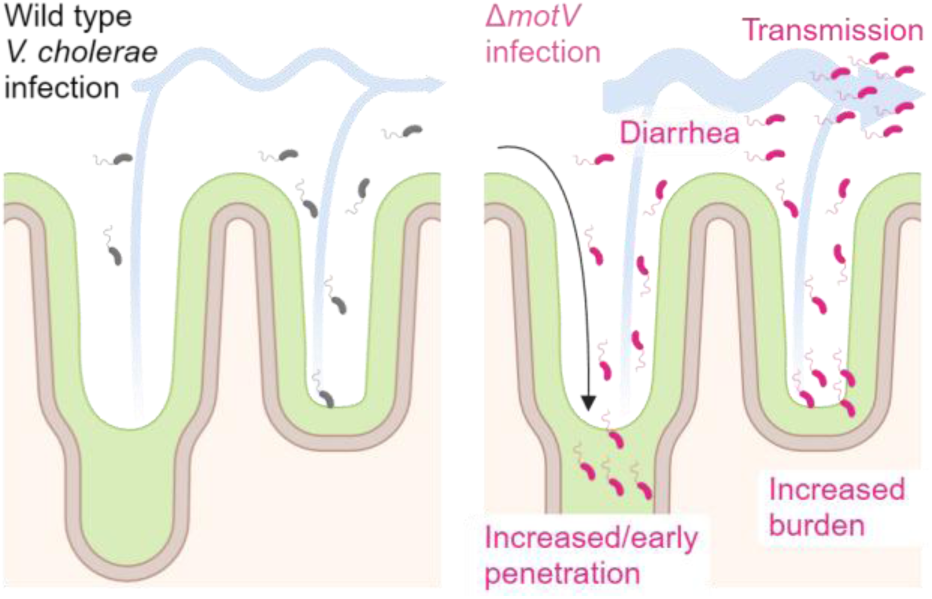
Model connecting the heightened inter-animal transmission of Δ*motV V. cholerae* to aberrant motility, bottleneck evasion, increased burden, and increased diarrheaogenicity. We propose that the *ΔmotV* mutant’s constitutive and rapid motility enables *V. cholerae* to penetrate deeper into the more proximal crypts of the SI early during infection and that this aberrant pattern of localization accounts for its hyper-transmissibility. The expansion of the *V. cholerae* intestinal niche permits more of the inoculum to survive the colonization bottleneck and replicate, increasing the burden of *V. cholerae* in the intestine and leading to greater diarrhea. Greater burden and diarrheal flux facilitate the excretion of the pathogen into the environment, promoting transmission.

We observed that the Δ*motV* mutant had an unprecedented property – it largely escaped the host infection bottleneck (Fig 4). In general, we interpret bottlenecks as host barriers to infection, and that host and/or microbiota changes can modulate the bottleneck. For example, the low pH of the stomach and the presence of an intact microbiota both contribute to the bottleneck restricting enteric colonization by the murine pathogen *Citrobacter rodentium* ^36,37^. The *motV* mutant is a new example of an emerging paradigm ^38,39^ that pathogen factors contribute to the bottleneck.

We propose that the circumvention of the bottleneck by the Δ*motV* mutant is a consequence of its aberrant swimming behavior and greater localization to the SI crypt niche. Paradoxically, by this logic, MotV activity in wild-type *V. cholerae* subjects the pathogen to more stringent population restriction. This may reveal that a major underlying contributor to the *V. cholerae* bottleneck in CD1 infant mice is the removal of bacteria from the SI by intestinal peristalsis.

By escaping the colonization bottleneck, bacteria lacking *motV* remain at a higher burden throughout the small intestine for the entire infection. The increased colonization of Δ*motV V. cholerae*, particularly in the proximal SI, is similar to that observed with *cheY* chemotactic mutants (Fig 2A-C) ^15,16,18^. Although *motV* is not known to be linked to the output of *V. cholerae’s* complex chemoreceptors and chemosensory pathways, deletion of *motV* and *cheY* cause similar motility phenotypes ^27^. Cells lacking *motV* or *cheY* are both only capable of counterclockwise flagellar rotation; they are unable to reverse the direction of flagellar rotation from counterclockwise to clockwise and consequently show increased smooth swimming in straight lines, decreased tumbling, and are unable to swim in soft agar ^18,27^. Studies using fluorescently labeled chemotactic deficient *V. cholerae* suggested that chemosensory pathways are not required for *V. cholerae* to penetrate into the intestinal crypts or for the localization of the pathogen along the crypt-villus axis ^29^. The similar swimming behavior and colonization phenotypes of the *cheY* and *motV* mutants suggest that their shared capacity for heightened colonization of the proximal SI may be caused by their constitutive straight swimming phenotype.

We propose that the aberrant swimming behavior and greater localization of the Δ*motV* mutant to the small intestine crypts increases disease in infected hosts. The swimming behavior of Δ*motV* may increase diarrheaogenicity by bringing the pathogen into a virulence-stimulating location and/or increasing the efficacy of toxin delivery by locating the pathogen closer to the epithelium. Moreover, the increased pathogen burden in animals infected with the Δ*motV* mutant likely also increases the quantity of toxin production, although elevated pathogen burden alone cannot fully explain the heightened diarrhea as *cheY* mutants have similar increased colonization without increasing diarrhea. While the molecular mechanism linking *motV* to intoxication remains to be described, the difference in diarrheaogenicity of the *cheY* and *motV* mutants provided a tool to probe the role of toxigenic diarrhea in *V. cholerae* transmission.

The Δ*motV* mutant’s increased colonization, virulence, evasiveness, and transmission could all be linked to its aberrant swimming behavior. Although motility is important in *V. cholerae* virulence, many pathogenic bacteria have lost their flagellum (e.g., *Shigella spp., Mycobacterium tuberculosis, Klebsiella pneumoniae*) or suppress expression of their flagellum during infection (e.g., *Listeria monocytogenes) ^40–44^*. As bacterial flagella are highly immunostimulatory, the lack of flagella in some pathogenic bacteria is generally believed to be an evolved mechanism for immune evasion. *V. cholerae’s* extreme motility during infection ^29^ suggests that motility provides an evolutionary advantage to this pathogen that outweighs the downsides of stimulating the immune system. By linking increased motility to transmission, the phenotypes of the Δ*motV* mutant highlight the potential evolutionary advantages of pathogen flagellation during enteric infection.

### Limitations of this study

The evolutionary benefits of *motV* are unclear. Presumably, *V. cholerae* motility and chemotaxis promote survival in the pathogen’s natural environments. By contrast, we find that MotV’s response to the stimuli encountered inside of the infant mouse SI is apparently maladaptive to survival/growth; the absence of *motV* promotes *V. cholerae* intestinal colonization and transmission between suckling mice, properties that seem beneficial for the propagation of the pathogen. These findings suggest a limitation of this otherwise valuable model. Possibly, the increased disease associated with the deletion of *motV* may increase mortality and be evolutionarily disadvantageous in humans, *V. cholerae’s* natural host. However, regardless of the evolutionary logic for a gene like *motV*, our findings with the Δ*motV* mutant illustrate the connection between intra-animal pathogenesis and inter-animal transmission.

The principal focus of this study was the characterization of the many infection-related phenotypes of a *V. cholerae motV* mutant, and how these phenotypes contribute to inter-animal transmission. Strains lacking *cheY* activity have previously been extensively characterized ^14–16,18,45–47^. The differences between *cheY* and *motV* mutants are that only the deletion of *motV* increased virulence and inter-animal transmission. The molecular mechanisms accounting for the different infection phenotypes of these stains may result from differences in their respective intra-intestinal localization, virulence regulation, and/or colonization dynamics and represent an interesting area for future study. Regardless, the divergence of phenotypes between the *motV* and *cheY* mutants provides a valuable tool for dissecting the mechanisms of *V. cholerae* transmission.

## Supplemental Figures

**Supplemental figure 1.**
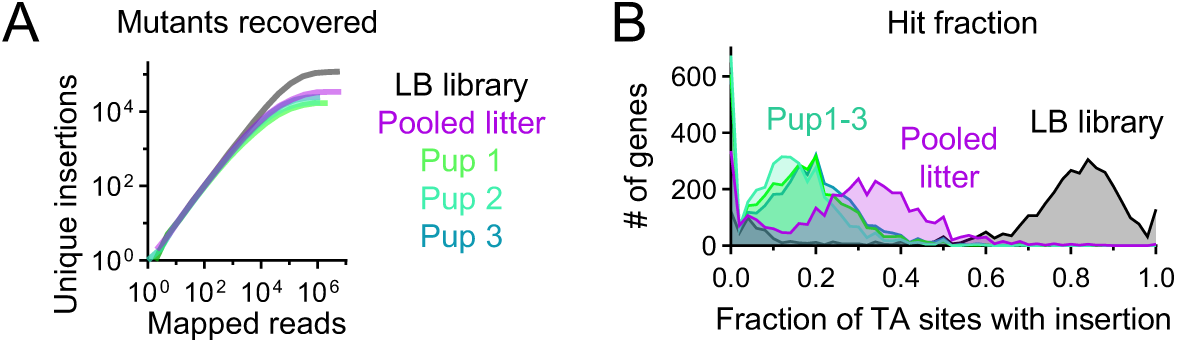
Pooling a litter of CD1 pups increases the fraction of the transposon library recovered after infection. Extended data from Fig 1. P4 CD1 mice were intragastrically inoculated with ∼5x10^7^ CFU of a mariner transposon library. 18-hours later, *V. cholerae* from the SIs of individual pups or the pooled SIs of an entire litter (10 pups) were outgrown overnight on LB, and then sequenced. For comparison, the same library was passaged overnight on LB media (LB library). **A,** The reads from each library were randomly sampled, and the number of unique insertion sites were plotted to depict diversity across sampling depths. **B,** Histogram of fraction of possible insertion sites in each gene disrupted by a transposon insertion. TA sites are sites with a TA dinucleotide. Mariner transposons primarily integrate at TA sites, and data throughout the manuscript is filtered only to include transposons integrated at TA sites.

**Supplemental figure 2.**
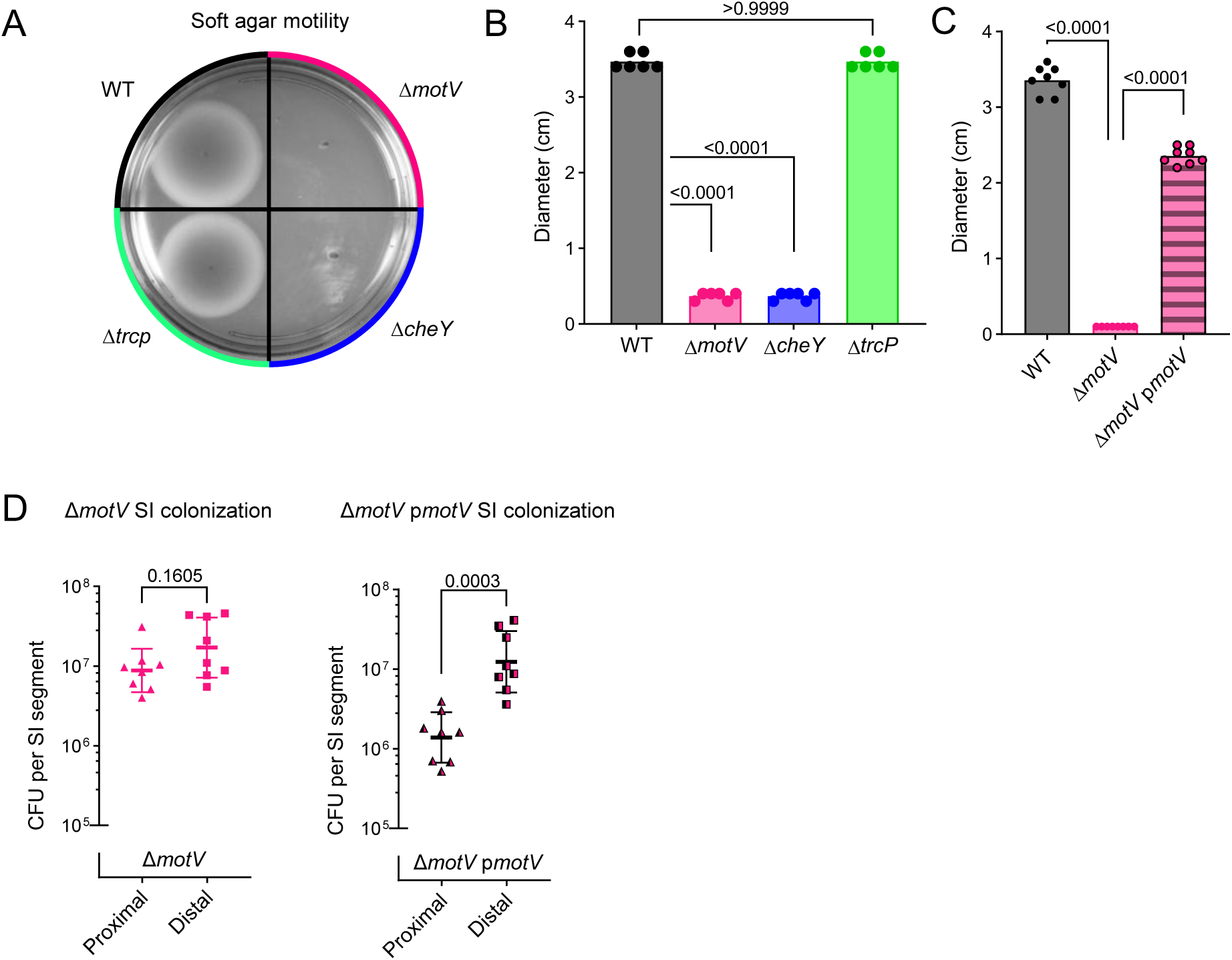
Extended observations and complementation of the Δ*motV* mutant. Extended data related to Fig 2. **A-C,** Swarming motility of indicated *V. cholerae* strains in soft agar plates. **A,** Image of a representative plate. **B-C,** Diameter of motility zones. A Δ*trcp* mutant is included as an additional control. Complementation of the Δ*motV* mutant with a plasmid-borne copy of *motV* (p*motV*) restores motility in soft agar. Mean and individual replicates. One-way ANOVA with Dunnett’s multiple comparison correction. **D,** CD1 mice were intragastrically inoculated with the Δ*motV* mutant or Δ*motV* complemented with p*motV*, and 18-hours later CFU was enumerated from the indicated SI segments. Geometric mean and standard deviation. Mann-Whitney test.

**Supplemental figure 3.**
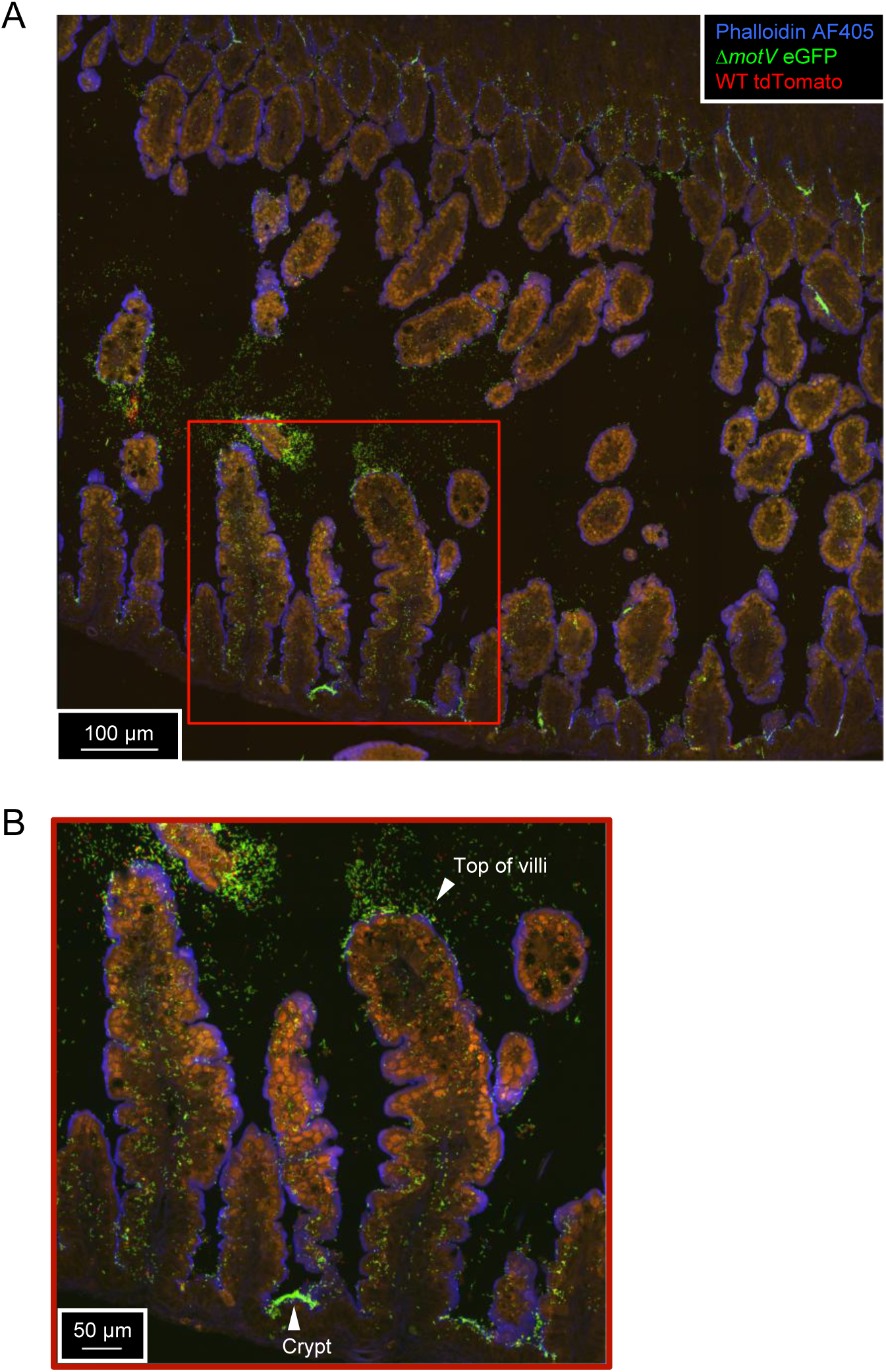
Large image of *V. cholerae* localization in the SI. Related to Fig 2. **A,** Large image of a 10 µm cryosection of the medial SI 18-hours after inoculation with ∼2 x 10^6^ CFU of a 1:1 ratio of WT *lacZ::tdTomato V. cholereae* (red) and Δ*motV lacZ::eGFP V. cholerae* (green). Shown are the villi of the SI on either side of the lumen. Sections were stained with AF405-conjugated phalloidin (blue) and imaged on a spinning disk confocal microscope. **B,** Zoomed-in portion of **A** (400 x 400 µm) showing the top of villi and crypts of the SI in more detail; outlined in red on **A**.

**Supplemental figure 4.**
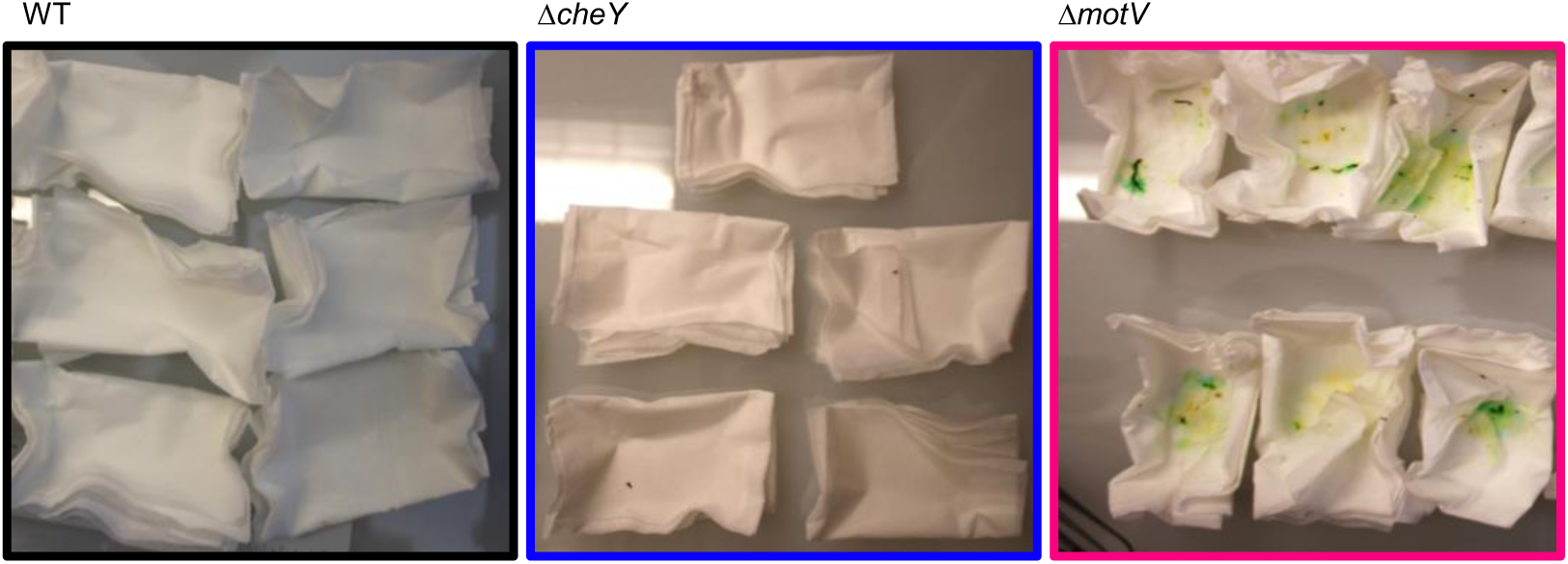
Deletion of *motV* increased infant mouse diarrheal discharge during infection with *V. cholerae*. CD1 pups were intragastrically inoculated with the indicated *V. cholerae* strains and individually housed for 18-hours in the pictured bedding. Each tissue is the bedding of a single infected animal. The *V. cholerae* inoculum is dyed green to assist with intragastric gavage, and subsequent discharge of the dye stains the bedding, assisting in visualizing diarrhea discharge.

**Supplemental figure 5.**
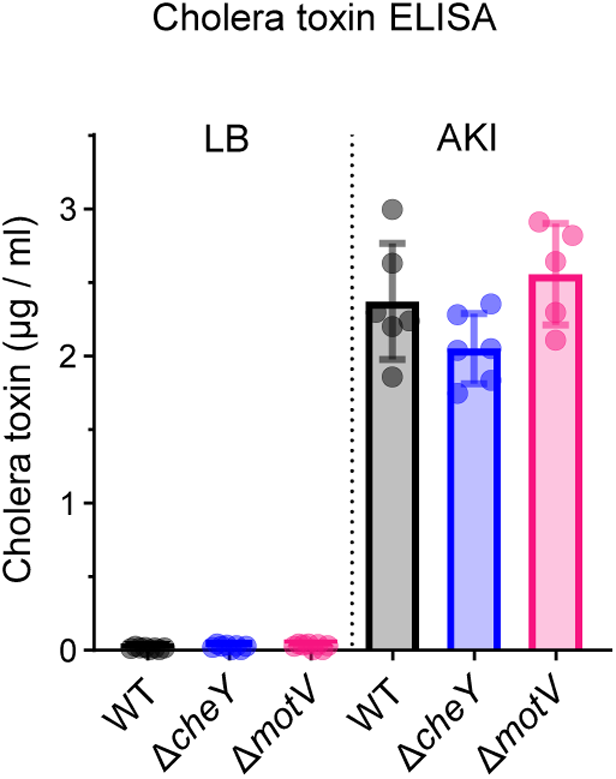
*motV* deletion does not change cholera toxin production in culture. ELISA measurement of cholera toxin following culture. Mean and standard deviation. Not significant by one-way ANOVA with Turkey’s multiple comparison correction.

**Supplemental figure 6.**
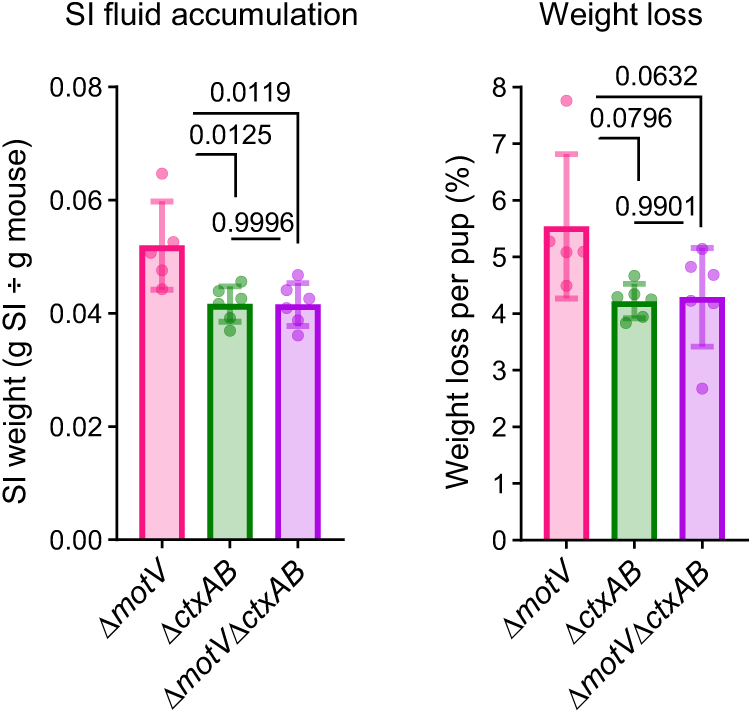
Deletion of *ctxAB* decreases the diarrheaogenicity of the Δ*motV* mutant. CD1 pups were intragastrically inoculated with the indicated *V. cholerae* strains and individually housed for 18-hours. Fluid accumulation in the SI was measured by comparing the weight of the animal to the weight of the SI. Weight loss was determined by comparing animal weight before and after infection. Mean and standard deviation. One-way ANOVA with Turkey’s multiple comparison correction.

## Methods

### Bacterial strains and growth conditions

Unless otherwise noted, bacteria were grown at 37 °C in lysogeny broth (LB) in liquid culture shaking at 200 rpm or on solid media containing 1.5% agar (weight/vol.). For the induction of cholera toxin, *V. cholerae* was grown in AKI broth (0.5% sodium chloride, 0.3% sodium bicarbonate, 0.4% yeast extract, and 1.5% Bacto peptone) for 4-hours static, followed by 4-hours with rotation at 37 °C ^48,49^. All *V. cholerae* were derivatives of a spontaneous streptomycin (Sm) resistant mutant of a 2010 *V. cholerae* clinical isolate from Haiti ^20^. Selection was accomplished by supplementing media with 200 µg/mL Sm. If required, antibiotics or other supplements were used in the following concentrations: carbenicillin (Cb), 100 µg/mL; kanamycin (Km), 50 µg/mL; diaminopimelic acid (DAP), 0.3 mM; sucrose (Suc), 0.2%; 5-bromo-4-chloro-3-indolyl-β-d-galactopyranoside (X-gal), 60 µg/mL. Bacterial stocks were stored at -80 °C in LB containing 25-35% (vol./vol.) glycerol. Strains used in this study are listed in Table S2.

### Strain and plasmid construction

*V. cholerae* strains with in-frame deletions were constructed using the vector pCVD442 and allelic exchange ^50^. 500-1,000 bp homology arms upstream and downstream of the target open reading frame were cloned into pCVD442 and introduced into *V. cholerae* by conjugation. Integration of the non-replicative plasmid was selected on Sm+Cb and double cross-overs removing the antibiotic-resistance cassette was selected for by growth in sucrose. Deletions were confirmed by PCR. WT *V. cholerae lacZ::tdTomato* was previously described ^15^, and the same method and vector (pJZ111) was use to introduce *eGFP* into the *lacZ* loci of the Δ*motV* mutant to create Δ*motV lacZ::eGFP*. p*motV* was constructed by integrating the *motV* open reading into the pBR322 vector ^51^ downstream of the P2 promoter.

### Mice

Animal studies were conducted at the Brigham and Women’s Hospital in compliance with the Guide for the Care and Use of Laboratory Animals and according to protocols reviewed and approved by the Brigham and Women’s Hospital Institutional Animal Care and Use Committee (protocol 2016N000416).

CDI dams with litters (mixed sex) were purchased from Charles River Laboratories (strain #022). Mice were housed in a biosafety level 2 (BSL2) facility under specific pathogen free conditions at 68-75 °F, with 30-50% humidity, and a 12 h light/dark cycle. CD-1 infant mice at postnatal day 3-5 were intragastrically inoculated *^30^* with the indicated dose and strain of *V. cholerae* in 50 µL of LB. Except for survival and transmission experiments, the infant mice were housed separately from the dams in tissue-lined boxes during infection. For survival and transmission experiments, pups were returned to dams where they remained until morbidity or the predetermined end point of the experiment. Infant mice were euthanized by isoflurane inhalation followed by decapitation. Adult mice were euthanized at the end of the study by isoflurane inhalation followed by cervical dislocation.

### Infant mouse colonization

For colonization assays, infant mice were separated from their dams and intragastrically inoculated with the indicated dose and strain. Dose (colony forming units; CFU) was determined by retrospective serial dilution and plating on selective media containing Sm. At the indicated times (<24-hours post inoculation), pups were euthanized, and the SI was removed. Colonization burden was determined in either the whole SI, or the SI divided into 2 or 3 parts of equal length (proximal, medial, and distal). Tissue segments were further divided in half for experiments that combined measurements of bacterial localization (microscopy) and bacterial burden. To disassociate the bacteria from the tissue, organs were homogenized in LB using a bead beater (BioSpec Product, Inc) and 2 stainless-steel 3.2 mm beads. SI CFU was determined by serial dilution and plating on selective media containing Sm.

Competitive infections were performed the same as other colonization experiments, except that the inoculum was a ∼1:1 mixture of the indicated strains. Blue/white CFU were determined by plating organ homogenates on LB + Sm/X-gal. Competetitive index was calculated by dividing the blue:white ratio from the SI by the blue:white ratio from the inoculum.

To measure diarrheaogenicity (diarrheal discharge, SI fluid accumulation, and weight loss), infant mice were individually housed for 18-hours following inoculation. The mice were weighed immediately following inoculation and immediately before euthanasia to determine weight loss. Following infection, the weight of the pup was compared to the weight of the entire resected SI to determine SI fluid accumulation. The bedding was weighed at the start and end of the experiment to determine diarrheal discharge.

### Tn-seq

Experiments were performed with a WT Himar transposon library ^52^. The library was expanded as a control on solid LB+Sm agar (LB library). For animal experiments, ∼5x10^7^ CFU of the library was inoculated into infant mice. 18-hours later, *V. cholerae* from the SIs of individual pups (Pup 1-3) or the pooled SIs of 10 mice from the same litter (Pooled litter) were outgrown overnight on LB+Sm agar. After outgrowth, cells were scraped off plates and frozen at -80 °C until processing.

Library preparation and sequencing were performed as previously described ^21,53^. gDNA was harvested using the GeneJet gDNA Isolation Kit. gDNA was fragmented to 400 bp using an ultrasonicator. Fragment ends were repaired with New England Biolabs Quick Blunting Kit and A-tailing was performed with Taq DNA-polymerase. Adaptor sequences were ligated onto fragments with T4 DNA ligase and used to perform PCR to enrich fragments containing the transposon. 300-500 bp fragments were isolated using gel extraction. Libraries were then sequenced on an Illumina NextSeq.

Sequencing data was processed using CLC Genomics Workbench (Qiagen) and the R-based RTISAn pipeline, as previously described ^21,54^. Trimming parameters: 3′ sequence – ACCACGAC; 5′ sequence – CAACCTGT; mismatch cost = 1; gap cost = 1; minimum internal score = 7; minimum end score = 4; discard reads <10 nt. Mapping parameters: reference – Haiti WT genome ^55^; mismatch cost = 1; insertion cost = 3; deletion cost = 3; length fraction = 0.95; similarity fraction = 0.95; reads mapped to multiple locations were mapped randomly; global alignment. RTISAn was used to convert mapping files to a positional tally (TAtally) exclusively counting insertions at TA dinucleotides. To account for the stochastic loss of mutants caused by infection bottlenecks, the LB library TAtally (input) was resampled 100 times to approximate the diversity of the animal TAtally (output). The simulations of the input were compared to the output to derive an average fold-change and *P* value.

To visualize the number of unique insertion sites per sample across a range of sampling depths, the TAtally was subjected to multinomial resampling beginning with the maximum read count of each sample and subsampling in two-fold lower increments. The number of unique insertions was calculated at each sampling depth. Prior to resampling, the TAtally was filtered by removing the bottom 1% of all reads to remove noise resulting from Illumina index sequence hopping.

### Microscopy

For microscopy, SI segments were retrieved as described for mouse colonization and then processed for cryosectioning ^56^. Tissue samples (∼ 2 cm) were fixed in PBS with 4% paraformaldehyde at 4 °C for 2-4 hours, placed in PBS with 30% sucrose overnight at 4 °C, and then mounted in a medium containing a 1:2.5 ratio of 30% Sucrose : Optical Tissue Clearing medium (OCT). Mounted tissue pieces were snap-frozen in liquid nitrogen, stored at -20 °C for 1-hour, then transferred to a -80 °C freezer until processing.

8-10 µm thick sections were cut on a Leica CM1860 cryostat and mounted on MAS adhesive slides (Matsunami Glass). Once samples were adhered to the slide, to preserve native fluorescence the samples were processed according to ^56^. After sections were cut, tissue pieces were delineated with a PAP-pen, then slides were dried in the dark for 20-minutes at room-temperature. Ice-cold 4% PFA was then added for 8-minutes while slides were stored in a humid chamber. PFA was removed and slides were then washed with PBS and 3% BSA-PBS. Then, a 1:1000 dilution of Alexa Fluor 405-conjugated phalloidin (ThermoFisher) was added to each segment for 15-minutes, in a dark humid chamber. After 15-minutes, slides were washed with PBS and 3% BSA-PBS again and mounted with VectaShield Plus non-hardening antifade mounting medium (vector laboratories). Slides were stored for 1-hour flat at room-temperature, then sealed with clear nail polish and stored at 4 °C until imaging. Slides were imaged with a Nikon Ti2 Eclipse spinning disk confocal microscope using a 40x oil immersion lens with a numerical aperture of 1.40 and an Andor Zyla 4.2 Plus sCMOS monochrome camera. Image analysis was performed on the ImageJ (FIJI (2.14.0)) software using custom macros.

For quantification of the localization of Δ*motV*:WT *V. cholerae* ratios in competitive infection, the number of WT (red) and Δ*motV* (green) *V. cholerae* were counted in 50-60 100x100 micron images per location (proximal/medial/distal; crypt/top of villus), taken from 20-30 larger (244.16x244.16 micron) fields. When 0 WT *V. cholerae* were observed, the Δ*motV*:WT ratio was set to 35 and labeled as “ND” (not detected), as the highest ratio of Δ*motV*:WT *V. cholerae* with WT organisms detected was 32:1.

For quantification of crypt occupancy, the total number of crypts in each of 20 244.16x244.16 micron fields per time point, infection, and location was recorded, as was the number of crypts in each field containing ≥1 *V. cholerae.* Crypt occupancy was determined by dividing the number of occupied crypts by the total number of crypts per field.

### Soft agar motility

1 µl of an overnight culture of the indicated strains were injected into semisolid LB agar (0.3%; weight/vol.). Plates were cultured at 37 °C for 6-hours and the colony diameter was measured. A WT control was included on every plate.

### Bottleneck quantification

Barcoded *V. cholerae* libraries were created with the plasmid donor library pSM1 ^57^. The pSM1 donor library contains ∼70,000 unique plasmids each with a random ∼25 bp barcode carried within a Tn7 site-specific transposon. Barcodes were introduced into the *V. cholerae* Tn7-integration site by triparental mating of the recipient *V. cholerae* strain with the pSM1 donor library and a helper plasmid (pJMP1039) that expresses the Tn7 transposase. Transconjugants were then selected on LB Sm Km plates. After outgrowth, transconjugates were harvested from selective plates in LB glycerol (25-35%; weight/vol.) and stored in aliquots at -80 °C.

STAMP libraries were inoculated and SIs were retrieved as described for other infant mouse colonization assays. *V. cholerae* burden (CFU) was quantified by serial dilution and plating on selective media. *V. cholerae* from the remaining sample was outgrown on LB Sm and bacterial samples were harvested and stored at -80 °C in LB glycerol (25-35%; weight/vol.) until processing. Samples were processed for barcode sequencing and STAMPR analysis as described in Hullahalli *et. al.,* 2021 *^31^*. Bacteria were boiled to release DNA and PCR was performed to amplify the barcode region and add sequencing adapters. Samples were sequenced on a NextSeq (Illumina). R and the STAMPR analysis pipeline ^31^ was used to demultiplex sequencing reads, trim, and map to the donor library pSM1. Founding population (Nr) was determined by STAMPR by comparing the number and frequency of barcodes recovered from the SI to a control sample outgrown from the animal inoculum. STAMPR scripts are available online at [https://github.com/hullahalli/stampr_rtisan].

### Infant mouse survival

Infant mouse challenge assays were performed as described in *Sit et. al.,* 2019 ^33^. Infant mice were orally inoculated with 10^6^ CFU of the indicated *V. cholerae* strain and returned to singly housed dams for maternal care. Survival was scored based on time from inoculation to morbidity. Body condition, diarrheal discharge, and temperature of infected mice were monitored every 4-6 hours to determine the onset of symptoms. Symptomatic mice were monitored every 30-minutes until reaching morbidity, at which point the pups were immediately removed and euthanized. At the predetermined endpoint (30-hours post inoculation), the remaining animals were scored as surviving and euthanized.

### Cholera toxin ELISA

*V. cholerae* strains were cultured in LB or AKI conditions. LB-culture: 8-hours in LB, with rotation, at 37 °C. AKI-culture ^48,49^: 4-hours static, followed by 4-hours with rotation at 37 °C in AKI broth (0.5% sodium chloride, 0.3% sodium bicarbonate, 0.4% yeast extract, and 1.5% Bacto peptone).

GM1 ELISA was used to quantify the concentration of cholera toxin in cell-free supernatant samples as described previously ^58^. Equal volumes of LB or AKI supernatants were serially diluted and known concentrations of purified cholera toxin were used as the standard. 96-well polystyrene microtiter plates were coated with GM1 ganglioside overnight, and 4 µg/ml fatty acid-free bovine serum albumin (BSA) was used to block the GM1-coated plates for 1 h at room temperature. Next, 260 μl of the supernatants were added to the wells in duplicate and incubated for 1 h at room temperature. Subsequently, a rabbit anti-CT polyclonal antibody (1:10,000) and then an HRP-linked goat ani-rabbit IgG antibody (1:5,000) were added to the wells and incubated for 1 h at room temperature each. For the development of the cholera toxin-antibody complex, tetramethylbenzidine (TMB) substrate solution (Thermo Fisher Scientific) was used according to the manufacturer’s protocol. The color intensity in each well was measured at 485 nm in a plate reader. The absolute quantity of cholera toxin in the samples was estimated by comparison to the standard curve.

### Infant mouse transmission

Infant mice were randomly re-assorted to prevent litter bias. ∼1/3 of each litter were inoculated with the indicated *V. cholerae* strain (seeds) and returned to foster-dams for maternal care with naïve littermates (contacts). Seeds and contacts remained with dams for 20-hours, at which point seeds and contacts were removed and euthanized. Transmission to contacts was determined by enumerating CFU in the SI. Contacts with 0 CFU in the SI were determined to be uninfected and contacts with ≥1 CFU were determined to be infected.

### Software and statistics

Data analysis was performed using CLC Genomics Workbench, R, GraphPad Prism, and Excel. Information regarding the number of samples and statistical tests are described in the figure legends. Geometric means, geometric standard deviations, and non-parametric tests were used for analyzing Tn-seq data, bacterial burden, bacterial competition, crypt occupancy, and founding population. Means, standard deviations, and parametric tests were used for comparisons of bacterial motility, diarrheal discharge, SI fluid accumulation, animal weight loss, and cholera toxin. Graphics and figures were prepared with BioRender, GraphPad Prism, and PowerPoint.

## Acknowledgements

We thank members of the Waldor lab for helpful discussions and feedback on the manuscript.

## License Information

This article is subject to HHMI’s Open Access to Publications policy. HHMI lab heads have previously granted a nonexclusive CC BY 4.0 license to the public and a sublicensable license to HHMI in their research articles. Pursuant to those licenses, the author-accepted manuscript of this article can be made freely available under a CC BY 4.0 license immediately upon publication.

